# Quantifying and interpreting the inverse coefficient of variation from minirhizotron-derived root dynamics

**DOI:** 10.64898/2026.11.24.768854

**Authors:** Julia Mayr, Yann Hautier, Alexandra Weigelt, Gorge A. Kowalchuk, Thomas Niemeyer, Vicky M. Temperton, M. Trinidad Torres-García, Emanuela W.A. Weidlich, Benjamin M. Delory, Kathryn E. Barry

## Abstract

1. The inverse coefficient of variation (μ/σ) is widely used to quantify temporal invariability of ecosystem functioning. However, most studies have focused on aboveground biomass production, while belowground processes remain understudied. Minirhizotrons enable repeated observations of root dynamics, but the resulting data differ fundamentally in their temporal structure from measurements obtained through aboveground biomass harvests, sequential soil coring, or ingrowth cores. This raises questions about whether inverse CV estimates depend on the metric used to represent root dynamics and on when and how frequently observations are made.
2. Using a five-year minirhizotron dataset, we quantified the inverse CV from standing root abundance and two change-based metrics (net change and rate of change in standing root abundance). Standing root abundance was quantified from individual imaging sessions, whereas change-based metrics were derived from differences between successive sessions. We compared their inverse CV estimates using all 45 imaging sessions, monthly observations (n = 35), and annual observations (n = 5). For annual observations, we compared measurements taken before the aboveground biomass harvest and at peak root abundance.
3. Inverse CV estimates differed between standing root abundance and change-based metrics, and correlations were weak and often negative. Estimates also differed across measurement frequencies and timings, with responses depending on the underlying metric. Together, these results show that inverse CV estimates depend on how root dynamics are represented and sampled.
4. We propose that the inverse CV does not always represent invariability and that its interpretation depends on the metric used to represent root dynamics. For standing root abundance, it reflects pool invariability. For change-based metrics, we suggest interpreting it as directional consistency of root gains or losses. This distinction may extend to other ecosystem fluxes that can be positive and negative, such as carbon or energy fluxes.
5. *Synthesis*. Our findings challenge the assumption that applying the same formula (μ/σ) ensures comparability and emphasize the need to align the underlying ecological questions and quantities when comparing invariability across ecosystem realms or studies.

## Introduction

Global environmental change is altering ecosystems worldwide, affecting their capacity to provide key functions and services such as biomass production, nutrient cycling, and carbon storage (Allan et al., 2015; Felipe-Lucia et al., 2020; IPCC, 2019). Understanding how these ecosystem functions vary through time is therefore a central objective in ecology.

Temporal variation in ecosystem functioning is commonly quantified using the coefficient of variation (*CV = σ/μ*) or its inverse (*inverse CV = μ/σ*), which are often interpreted as measures of temporal variability and invariability, respectively (e.g., Isbell et al., 2026; S. Wang et al., 2019). Empirical studies have predominantly relied on the inverse CV of plant aboveground biomass production as a measure of ecosystem invariability. This emphasis likely overlooks the substantial contribution that belowground processes can make to ecosystem functioning and its temporal variation (Bardgett & van der Putten, 2014; Z. Zhang et al., 2026).

In grasslands, roots can account for more than half of total plant biomass and often exhibit different temporal dynamics and responses to environmental fluctuations (Bonato Asato et al., 2025; Dohleman et al., 2012; Ma et al., 2021). Furthermore, roots enable persistence during and recovery after stress in two ways. First, roots acquire water and nutrients, supported by exudation and symbiotic associations that mobilize otherwise inaccessible soil resources (Holz et al., 2024; Teotia et al., 2017). Plasticity in root system architecture and trait combinations further allow roots to maintain resource acquisition under changing environmental conditions (Li et al., 2017; Viana et al., 2022). Second, roots remain viable during stressful conditions, while structures like storage organs and bud banks allow shoot regrowth following disturbance (Ott et al., 2019; Tumber-Dávila et al., 2026).

However, understanding how roots contribute to variation in ecosystem functioning remains methodologically challenging because root systems are inaccessible and sensitive to disturbance (Freschet et al., 2021). Root biomass is typically estimated in two ways. First, soil cores can be extracted, sometimes sequentially, to estimate root biomass at different soil depths. This method is simple, scalable, and cost-effective, but provides only snapshots in time, offers no direct measure of production or turnover, and often yields noisy estimates due to high spatial variability and difficulties in separating live from dead roots (Freschet et al., 2021; Milchunas, 2009). Second, ingrowth cores use a defined volume of root-free soil into which roots grow over a specific time interval to estimate production. However, root growth into ingrowth cores can be biased, with evidence for both root avoidance and artificially stimulated proliferation depending on mesh size, fill material, bulk density, and incubation time (Addo-Danso et al., 2016; Freschet et al., 2021; Milchunas, 2009). Although widely used, both approaches are destructive, labor-intensive, and limited in capturing continuous root dynamics over time.

Minirhizotrons offer a promising alternative to destructive sampling techniques. They involve the initial installation of transparent observation tubes into the soil, allowing researchers to repeatedly image the same area of soil through time. After installation, this technique enables high-resolution, non-destructive observations of root birth, growth, and lifespan across the soil profile under field conditions (Addo-Danso et al., 2016; Milchunas, 2012). As with any field-based technique, minirhizotrons are subject to methodological constraints, including preferential root growth along the soil-tube interface, a limited field of view, and labor-intensive image processing workflows (Addo-Danso et al., 2016; Freschet et al., 2021; Parker et al., 1991; Wahlström et al., 2015). Minirhizotrons may, however, be particularly well suited to detect temporal variation that is difficult to observe using destructive methods (Addo-Danso et al., 2016). Recent advances in deep learning have further improved their efficiency by accelerating root segmentation, reducing observer bias, and improving reproducibility in root detection (Peters et al., 2023; Smith et al., 2022).

Minirhizotrons offer a unique opportunity to study temporal variation in root dynamics across years, within seasons, and across soil depths. However, three analytical challenges remain. First, measurement frequency may influence inverse CV estimates because the temporal variation captured depends on how frequently roots are observed (Stewart & Frank, 2008; Tingey et al., 2003). Moreover, because measurement frequency determines the number of observations in a time series of fixed duration, inverse CV estimates may vary as a function of measurement frequency alone (Chao et al., 2025). Second, measurement timing may influence inverse CV estimates because the timing of root growth and mortality varies throughout the year (Bonato Asato et al., 2025; Radville et al., 2016). Measurements collected at different times of the year may characterize different phases of this seasonal cycle. Third, root dynamics from minirhizotron images can be quantified using either standing root abundance proxies such as planar root length density (*pRLD*; Wacker et al., 2025) from a single image, or difference-based “productivity” metrics from sequential images. Accordingly, the same minirhizotron observations can be used to quantify root dynamics using metrics that approximate either a pool or a flux, and neither is a simple analog of how we measure aboveground biomass production to quantify the inverse CV aboveground (Text Box 1).

Here, we explore how the inverse CV derived from minirhizotron images depends on these three analytical decisions. We examine how (i) measurement frequency, (ii) measurement timing, and (iii) metric choice (abundance vs. change) influence resulting inverse CV estimates. We further evaluate how these decisions affect the ecological interpretation of the inverse CV and aim to provide a foundation for future studies using minirhizotrons to investigate temporal variation in root dynamics.

## 1. Materials and Methods

### 1.1. Data source and minirhizotron time series

We used minirhizotron data from the Priority Effects Mechanisms (POEM) field experiment in northern Germany, an experimental platform designed to test how priority and year effects influence community assembly and ecosystem functioning (Alonso-Crespo et al., 2025). POEM consists of multiple sub-experiments established in different years where the order of arrival of three plant functional groups (grasses, forbs, and legumes) was manipulated (see Alonso-Crespo et al., (2025) for a full description of the POEM experiment). We focused on the sub-experiment established in 2021 (POEM2021), which provides a multi-year record of root dynamics. POEM2021 consists of 25 mixture plots (each 3 m x 3 m), of which 20 were sown with experimentally assembled grassland communities differing in plant functional group order of arrival, while five were left unsown to allow free succession.

We installed two 1 m long and 60 mm wide acrylic tubes at a 45° angle in each of the 20 sown mixture plots six months before the start of POEM2021, resulting in 40 tubes in total. We use data from these 40 tubes as a case study to evaluate how methodological decisions influence inverse CV estimates derived from minirhizotron observations, independent of treatment effects. Each tube was imaged at 18 evenly spaced soil depths between 1.4 and 49.5 cm deep. Images were collected using a VSI MS-190 manual minirhizotron camera (Vienna Scientific Instruments GmbH) at a resolution of 2340 × 2400 pixels, corresponding to 148 pixels mm⁻¹.

Root images were collected between April 2021 and November 2025 following a standardized imaging protocol, resulting in 45 imaging sessions. Images were collected twice a month from April–September 2021 and April–September 2022; once a month from October 2021–March 2022, October 2022–June 2023, April–August 2024, and from April–September 2025, with an additional session in November 2025. All images were processed using a two-step pipeline. First, we trained a convolutional neural network in RootPainter (Version: 0.2.28) for root segmentation (Smith et al., 2022). Second, we quantified total root length in segmented images using RhizoVision Explorer (*V2.0.3*; Seethepalli et al., 2021). By dividing the total root length of each image by the image area, we converted these measurements to planar root length density (*pRLD*; cm cm⁻²), which we used as our metric of standing root abundance for all subsequent analyses.

### 1.2. Time series under different measurement frequencies

To evaluate how measurement frequency affects inverse CV estimates, we derived three tube-level *pRLD* time series with different numbers of observations. First, time series based on all measurements included all available imaging sessions from April 2021 to November 2025 (45 time points), thereby retaining the full temporal variation captured by the minirhizotron observations (Fig. 1). During 2021–2022, roots were imaged approximately twice per month during the growing season. This period coincides with the highest rates of root production and turnover in temperate grasslands (Steinaker & Wilson, 2008).

**Fig. 1.**
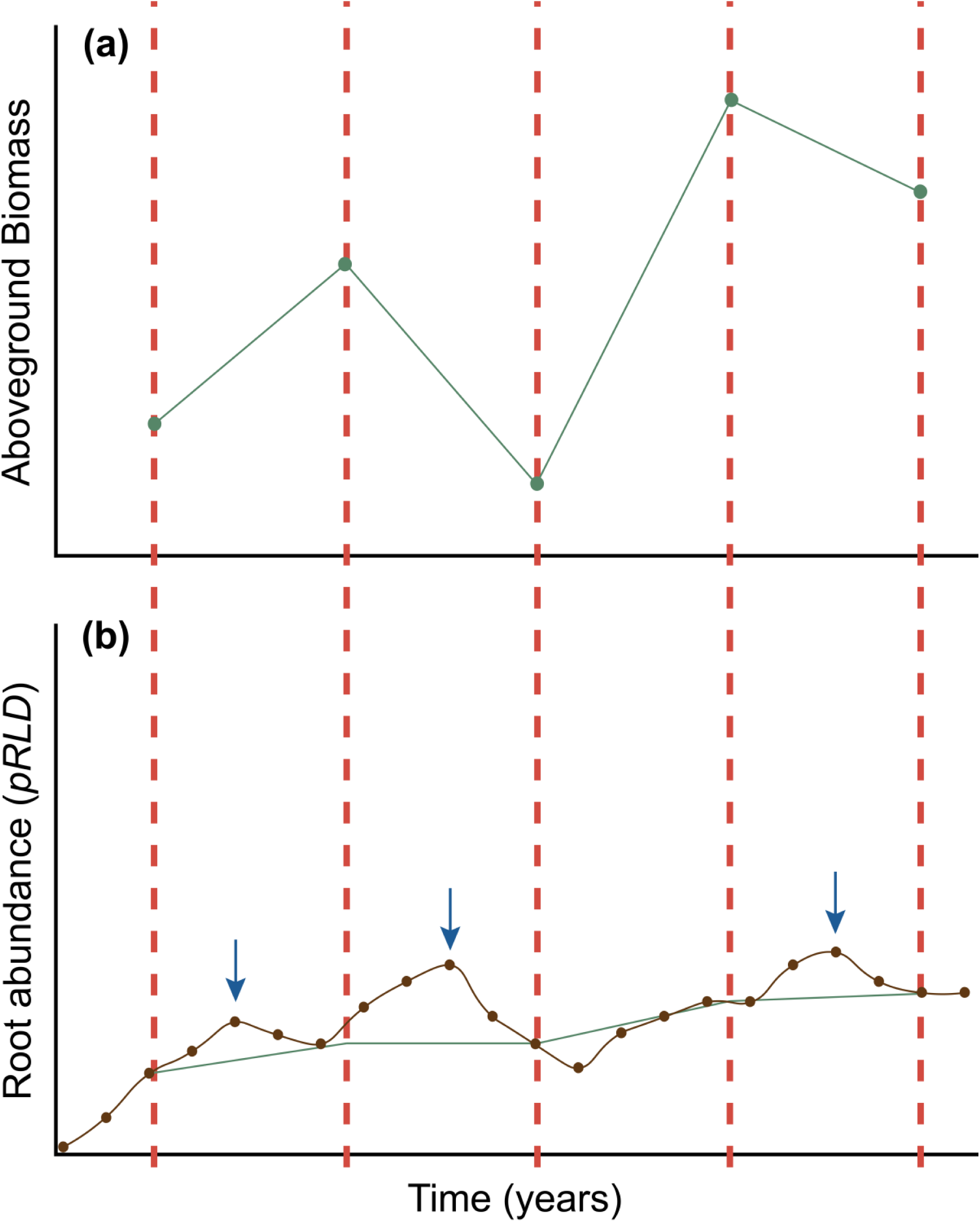
Conceptual illustration of aboveground biomass and minirhizotron-derived root abundance (*pRLD*) over time. (a) Aboveground biomass is typically harvested once per year at a fixed time (red dashed lines), yielding a single annual measurement. (b) Minirhizotron-derived *pRLD* captures fine-scale temporal variation (brown line). Sampling root abundance only once per year at the timing of aboveground biomass harvest (green line) produces a much coarser trajectory, which may not capture key root dynamics or peak abundance (blue arrows).

Second, for the monthly measurements, we selected one imaging session per calendar month. When multiple sessions occurred within a month, we chose the session that produced the most even spacing relative to the preceding and following sessions. This measurement frequency resulted in 35 time points and reflects the coarser sampling intervals present in parts of the dataset.

Third, for the annual measurements, we selected a single imaging session per calendar year, resulting in 5 time points. This measurement frequency corresponds to the annual aboveground biomass measurement typically used to quantify temporal invariability. Because all time series were derived from the same underlying observations, differences among measurement frequencies could be assessed without confounding variation across years or sampling protocols.

### 1.3. Annual time series under different measurement timings

To evaluate the effect of measurement timing, we compared two annual sampling strategies, each selecting one imaging session per tube per year. First, we selected the imaging session most closely preceding the annual aboveground biomass harvest, which took place at the end of June or in early July (hereafter pre-harvest measurements). This calendar-based sampling strategy is most similar to how aboveground biomass is sampled in grassland studies of temporal invariability (Fig. 1). Second, for each tube and year, we identified the imaging session with the maximum observed *pRLD* (hereafter peak measurements). This phenology-based sampling strategy captures the maximum standing root abundance attained each year (Fig. 1). Although the selected imaging session may not coincide with the aboveground harvest, it is conceptually comparable because both aim to capture peak standing biomass.

### 1.4. Metric computation

For each imaging session, we averaged *pRLD* across all depths within a tube to produce a single value per tube. This aggregation allowed us to focus on methodological effects on inverse CV estimates at the tube level rather than on depth-specific variation. Within each time series, depth-averaged *pRLD* at a given time point served as a proxy for standing root abundance. We then calculated net change in root abundance as the difference in *pRLD* between consecutive imaging sessions (*ΔpRLD*). To account for unequal time intervals between sessions, we additionally calculated a time-standardized rate of change (*ΔpRLD / Δt*) by dividing the net change by the number of days between consecutive sessions. Both net change and rate of change can be positive or negative depending on the balance between root growth and mortality. As a supplementary analysis, we implemented a generalized additive model (GAM)-based workflow, in which the fitted model estimated standing root abundance, while its integral and derivative provided conceptually comparable measures of net change and the time-standardized rate of change (Appendix S2).

### 1.5. Calculating the inverse coefficient of variation

We quantified the inverse CV (μ/σ) for the three metrics (standing root abundance, net change, and rate of change) within each of the four tube-specific time series (all measurements, monthly measurements, annual pre-harvest measurements, and annual peak measurements). This resulted in 12 inverse CV estimates per tube. Inverse CV estimates for change-based metrics can be negative when their temporal mean is negative. Because measurement frequency determined how many observations were included in the time series, the number of observations underlying the inverse CV estimates differed. The coefficient of variation, and therefore the inverse CV, can be sensitive to the number of observations in the time series (Chao et al., 2025). Given this sensitivity, we used an information-based framework to standardize for unequal numbers of observations in a supplementary analysis. This framework requires strictly positive data; therefore, we applied it only to standing root abundance (Appendix S3).

### 1.6. Statistical analysis

To assess whether the inverse CV calculated from standing root abundance and from change-based metrics produced similar rankings of tubes, we calculated Spearman rank correlations between the abundance-based inverse CV and the inverse CV derived from net change and rate of change across the 40 minirhizotron tubes for each of the four time series.

We then fitted a hierarchical Bayesian distributional model to quantify how measurement frequency and timing, and metric choice, influence belowground inverse CV estimates while accounting for multiple estimates derived from the same minirhizotron tube. We implemented the model using the *brms* package (Bürkner, 2017) in R v.4.5.2 (R Core Team, 2025) with the *cmdstanr* backend (Gabry et al., 2025). The model was fitted using four Markov Chain Monte Carlo chains with 4,000 iterations each, including 1,000 warm-up iterations, and the default weakly informative priors.

We modeled inverse CV estimates as a function of measurement frequency and timing (treated as a single factor), metric, and their interaction. The model was specified without an intercept to allow direct comparison among all combinations of measurement frequency and timing, and metric. Residual variance was also allowed to vary across measurement frequency and timing, and metric. Because multiple inverse CV estimates were calculated per tube, we included tube as a hierarchical grouping variable to account for non-independence, using the simplest structure supported by model comparison (Appendix S1: Table S1). The model showed excellent convergence, with R-hat values of 1.00 for all parameters. Model performance and goodness-of-fit were evaluated using Leave-One-Out Cross-Validation and posterior predictive checks (Appendix S1).

## 2. Results

### 2.1. Comparison of inverse coefficient of variation estimates across metrics

Across the 40 tubes, Spearman rank correlations between abundance-and change-based inverse CV estimates were generally weak. Across all, monthly, and peak measurements, abundance-based inverse CV estimates were weakly negatively correlated with those based on both net change and rate of change (ρ = −0.25 to −0.16; Fig. 2). For pre-harvest measurements, correlations differed between change-based metrics. Abundance-based inverse CV estimates showed no clear relationship with those based on net change (ρ = 0.01) but were moderately negatively correlated with those based on rate of change (ρ = −0.46; Fig. 2). These correlations indicate low agreement in tube rankings, with tubes ranked differently depending on whether inverse CV estimates were based on abundance– or change-based metrics.

**Fig. 2.**
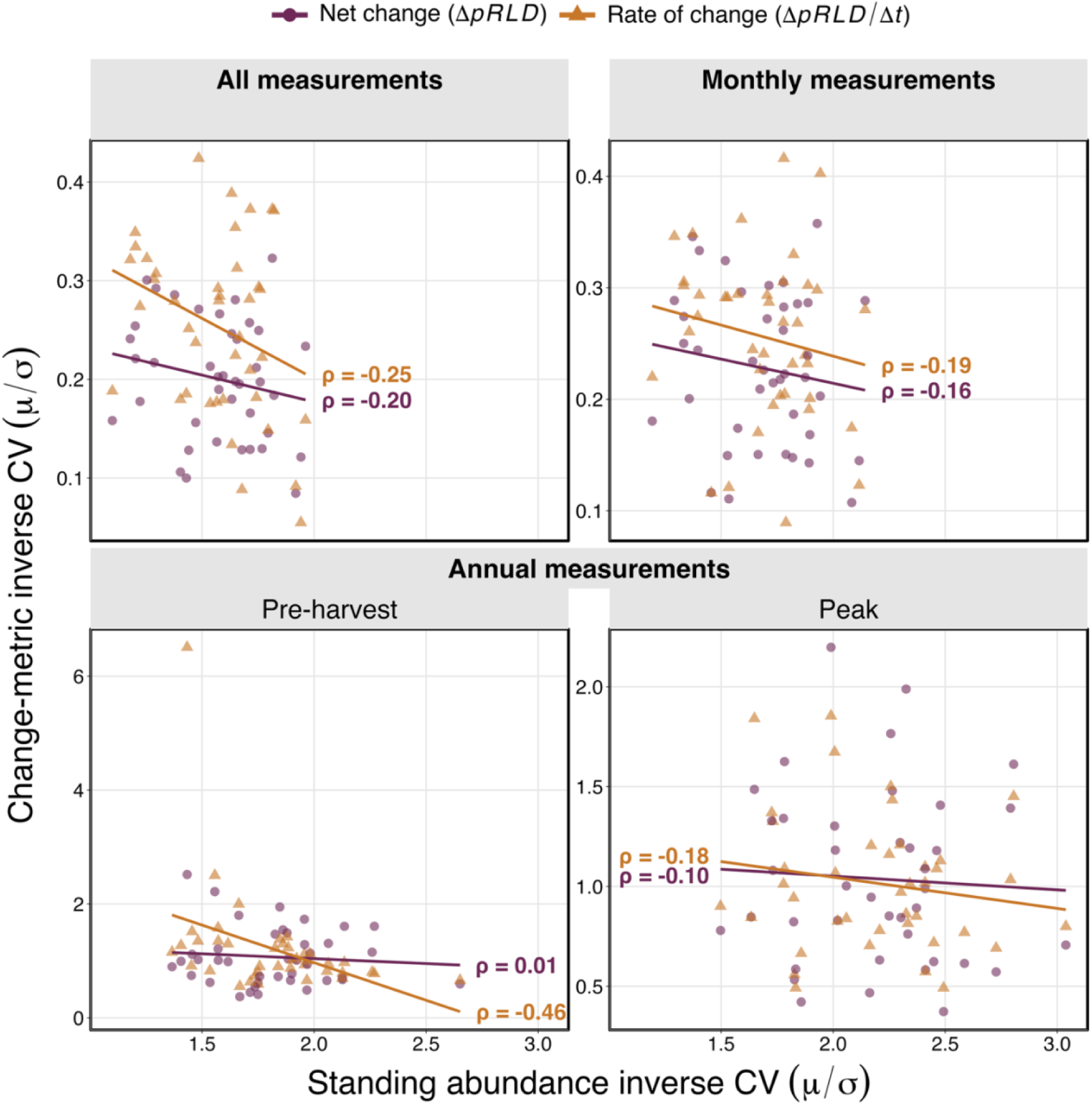
Relationships between abundance-based and change-based inverse CV (μ/σ) at different measurement frequencies and timings. Points represent tube-level inverse CV estimates (n = 40). Panels show all measurements, monthly measurements, and annual measurements at two timings (pre-harvest and peak). Lines show linear fits for visualization.

The hierarchical Bayesian distributional model also showed that, across measurement frequencies and timings, predicted inverse CV was higher for standing root abundance than for either change-based metric (Fig. 3; Appendix S1: Table S2).

**Fig. 3.**
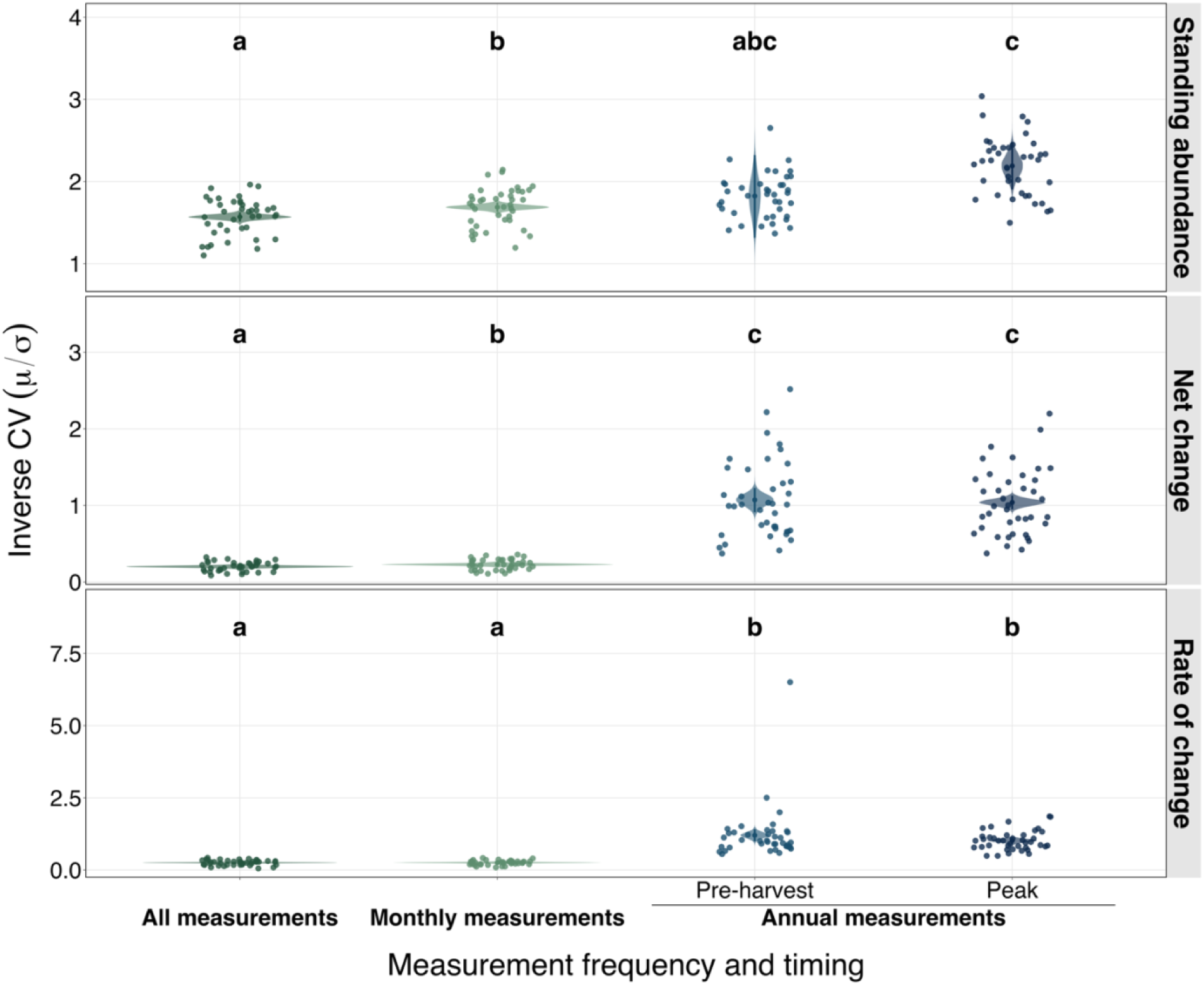
Effects of measurement frequency and timing, and metric on inverse CV estimates. Inverse CV (μ/σ) calculated from standing root abundance (*pRLD*), net change (*ΔpRLD*), and rate of change (*ΔpRLD / Δt*) for all, monthly, and annual measurements at two timings (pre-harvest and peak). Points show observed tube-level estimates (n = 40); model summaries indicate posterior means with 95% credible intervals. Letters summarize pairwise contrasts whose 95% credible intervals excluded zero (Appendix S1: Fig. S5).

### 2.2. Effects of measurement frequency and timing on inverse coefficient of variation estimates

Inverse CV estimates also varied with measurement frequency and timing within each metric (Fig. 3). For standing root abundance, posterior mean inverse CV increased with coarser measurement frequency and was highest for peak measurements (Fig. 3; Appendix S1: Table S2). All pairwise contrasts involving pre-harvest measurements had 95% credible intervals that overlapped zero (Fig. 3; Appendix S1: Fig. S5), indicating that pre-harvest measurements did not credibly differ from the other measurement frequencies and timings. When standing root abundance was analyzed using an information-based framework (Appendix S3), estimates increased with coarser measurement frequency and were higher for phenology-based than for calendar-based annual measurements (Appendix S3: Fig. S1 b,c), with 95% credible intervals for all pairwise contrasts excluding zero (Appendix S3: Fig. S1 b,d).

Change-based metrics (i.e., net and rate of change) showed similar responses across measurement frequencies (Fig. 3), with substantially higher inverse CV estimates for annual than finer measurement frequencies. For instance, posterior mean inverse CV was nearly five times greater for pre-harvest than for monthly measurements for both change-based metrics (Fig. 3; Appendix S1: Table S2). The two annual measurement timings did not credibly differ for either metric. Within the finer measurement frequencies, however, only net change showed a slight increase from all to monthly measurements (Fig. 3; Appendix S1: Fig. S5).

Abundance-based inverse CV had substantially greater variance than change-based inverse CV. Variance also increased with coarser measurement frequency and was highest for pre-harvest measurements, indicating greater differences among tubes (Appendix S1: Table S2).

When inverse CV was derived from GAM-smoothed root abundance trajectories rather than raw observations (Appendix S2), estimates increased with coarser measurement frequency across metrics. However, the effect of annual measurement timing differed among metrics.

Abundance-based inverse CV estimates were highest for peak measurements, whereas change-based inverse CV estimates were highest for pre-harvest measurements (Appendix S2: Fig. S1).

## 3. Discussion

The inverse coefficient of variation is widely used as a measure of temporal invariability in ecosystem functioning. However, empirical applications have focused predominantly on aboveground biomass production, while belowground processes remain understudied due to the inaccessibility of root systems and associated methodological constraints. Minirhizotrons enable repeated, high-resolution observations of root dynamics, but the resulting data differ fundamentally in their temporal structure from data on aboveground biomass typically used to quantify the inverse CV and from data obtained using commonly applied field methods (e.g., sequential soil coring or ingrowth cores). Quantifying the inverse CV from minirhizotrons therefore requires careful consideration of how root dynamics are represented, as well as when and how frequently they are measured. Using a multi-year dataset, we quantified the inverse CV from standing root abundance (our closest analog to aboveground standing biomass) and change-based metrics (our closest analog to productivity) across three measurement frequencies and two annual measurement timings. We found that inverse CV estimates differed between abundance– and change-based metrics and across measurement frequencies and timings, suggesting that inverse CV estimates depend on how root dynamics are measured and represented.

### 3.1. Inverse coefficient of variation estimates derived from pools and fluxes are not interchangeable

The same minirhizotron observations can be used to derive standing root abundance at each time point (e.g., *pRLD*), approximating a pool, or changes in root abundance between observations, approximating fluxes (Box 1). We therefore compared the inverse CV (μ/σ) derived from standing abundance– and change-based metrics. Inverse CV estimates calculated from standing root abundance were consistently higher than those calculated from change-based metrics. This difference partly arises because change-based metrics represent fluxes as increments between observations with temporal means often centered around zero, whereas abundance reflects a pool as a cumulative quantity with larger temporal means. Beyond differences in magnitude, correlations between abundance– and change-based inverse CV estimates were generally weak and often negative, indicating poor agreement in their relative ranking. As a result, any given tube could have a relatively high or low inverse CV depending on how root dynamics were represented, despite being derived from the same time series. Together, these results show that applying the same formula (μ/σ) to abundance– and change-based variables does not yield interchangeable inverse CV estimates, because pools and fluxes capture different aspects of temporal root dynamics.

The ecological interpretation of the inverse CV therefore depends on whether it is calculated from a pool, such as standing root abundance, or from a flux, such as change in root abundance between observations. When calculated from standing root abundance, inverse CV expresses **mean standing root abundance** relative to its temporal variability. It can therefore be interpreted as the consistency, or invariability, of a standing root pool through time. When calculated from change-based metrics, its interpretation shifts because losses can outpace gains over time, resulting in a negative mean and therefore a negative inverse CV. This complicates an interpretation in terms of invariability because the inverse CV can be positive or negative. In this case, inverse CV expresses **mean change in standing root abundance** relative to its temporal variability, with the signed mean indicating the prevailing direction of change toward gains or losses, and variability among increments contributing to how strongly that prevailing direction is reflected over time. For a given mean change, lower variability among increments therefore indicates greater consistency in the direction of change over time. We therefore interpret inverse CV calculated from change-based metrics as directional consistency, where the sign indicates the prevailing direction toward gains or losses, the absolute magnitude reflects the strength of the mean direction relative to variability among changes over time, and an inverse CV of zero indicates no prevailing direction of change. Importantly, directional consistency does not necessarily imply that individual changes have the same sign or that any direction persists sequentially through time. It also does not reflect the magnitude of fluctuations, as time series with larger or smaller fluctuations can have the same directional consistency (Appendix S4: Fig. S1).

Pool invariability and directional consistency are therefore not equivalent and do not necessarily follow the same pattern. For example, a system near equilibrium may have high pool invariability, with standing root abundance maintained around a stable level, and low directional consistency, with increments frequently alternating between gains and losses (Appendix S4: Fig. S2 a,b). Conversely, a system that grows continuously over the same period may have lower pool invariability because root abundance changes substantially over time, but higher directional consistency because net change is persistently positive and of similar magnitude (Appendix S4: Fig. S2 c,d). Disturbances that increase root mortality may modify both pool invariability and directional consistency, but through different changes in mean and variability (Appendix S4: Fig. S3), reinforcing that abundance– and change-based inverse CV estimates represent distinct but complementary dimensions of temporal variation.

### 3.2. Belowground pool and flux measures in the context of aboveground measures

The distinction between inverse CV calculated from pools and fluxes becomes particularly relevant when considering how it is quantified in practice. In grasslands, empirical studies have calculated the inverse CV from annual aboveground biomass harvests and interpreted it as temporal invariability of aboveground net primary productivity (ANPP; e.g., Hautier et al., 2015; Isbell et al., 2009). Although ANPP is conceptually defined as a flux representing biomass production over time (Roxburgh et al., 2005; Sala & Austin, 2000), it is often estimated from an annual harvest conducted near peak biomass (Sala & Austin, 2000). The quantity measured at harvest, however, more closely resembles a standing biomass pool. A key distinction from a true standing biomass pool is that the biomass is often removed at each harvest. The biomass measured at the subsequent harvest therefore represents regrowth over the intervening period, and time is implied. Furthermore, the harvested biomass does not capture losses that occur prior to harvest (e.g., herbivory, litterfall, decomposition, or senescence; Clark et al., 2001; Roxburgh et al., 2005). Consequently, annually harvested biomass does not correspond strictly to either a pool or a flux but combines characteristics of both. Accordingly, it differs from both standing root abundance and change-based metrics derived from minirhizotron observations in how temporal dynamics are represented and interpreted. In this respect, it is conceptually similar to root production estimated using ingrowth cores, which likewise reflects biomass accumulated over a defined interval and may not capture losses during that period (Katayama et al., 2019).

Studies of aboveground ecosystem functioning in forests distinguish more clearly between pool and flux variables. Standing biomass is quantified from repeated measurements of tree size (Chave et al., 2005; Clark et al., 2001), while growth over time is quantified as growth increments between censuses (Chambers et al., 2001; Clark et al., 2001). As a result, empirical studies using the inverse CV in forests also distinguish whether the underlying variable is a proxy for biomass (Yuan et al., 2019) or production (i.e., biomass gains; Qiao et al., 2023; R. Shan et al., 2025). For example, R. Shan et al., (2025) evaluated the relationship between forest stand age and invariability using inverse CV calculated from production (biomass gains only) and net change (gains and losses between consecutive censuses). These representations correspond closely to our distinction between standing root abundance as a proxy for a pool and changes in root abundance as proxies for fluxes. Consequently, the measures underlying minirhizotron-derived inverse CV estimates are more directly comparable to those used aboveground in forests than to annual harvest-based measures used in grasslands.

### 3.3. Measurement frequency and timing interact with metric choice

Measurement frequency also influenced inverse CV estimates derived from minirhizotron observations, with coarser frequencies generally leading to higher estimates. This effect likely arises from both statistical and biological mechanisms. First, statistically, CV (and its inverse) is sensitive to the number of observations in a time series (Chao et al., 2025), such that estimates can differ among measurement frequencies simply because they contain different numbers of observations. However, supplementary information-based analyses of standing root abundance (Appendix S3) showed the same pattern, with even clearer differences among measurement frequencies. Statistical mechanisms alone are therefore unlikely to explain this pattern. Second, biologically, high-frequency measurements allow short-term fluctuations in root growth and mortality within seasons to be resolved (Stewart & Frank, 2008; Tingey et al., 2003). At coarser measurement frequencies, short-term variation may be missed (e.g., Appendix S5: Fig. S1; Stewart & Frank, 2008), thereby contributing to higher inverse CV estimates.

The effect of measurement frequency and timing further depended on the metric used. For standing root abundance, inverse CV increased progressively from all to monthly and annual peak measurements, whereas annual pre-harvest measurements did not differ credibly from any other measurement frequency or timing. Root dynamics can vary substantially among years and are influenced by climatic conditions (Fiala et al., 2012; Hui & Jackson, 2006; J. Wang et al., 2019), such that calendar-based annual measurements at a fixed time may capture different phases of root growth or loss across years. These differences in phenological state captured across years may therefore add temporal variation in standing root abundance despite the lower measurement frequency. Phenology-based annual peak measurements instead align observations to a biologically comparable state across years, which may result in greater observed root pool invariability. At the same time, because annual peaks were identified retrospectively for each tube and year, the higher inverse CV may partly reflect the selection of observed maximum values rather than phenological alignment alone.

For change-based metrics, annual measurement timing had little effect. Inverse CV estimates did not differ between peak and pre-harvest measurements for either net change or rate of change. At annual measurement frequencies, both net change and rate of change reflect differences in root abundance between consecutive years, and the directional consistency of these year-to-year changes was similar whether measurements were aligned by phenology or calendar date. At higher measurement frequencies, however, the two change-based metrics showed different patterns. Inverse CV did not differ between all and monthly measurements for rate of change, whereas net change showed a small but credible difference. This likely reflects that net change depends on the interval length between observations, whereas rate of change is expressed as net change per unit time, reducing variation associated with temporal spacing and making inverse CV less sensitive to measurement frequency (e.g., Appendix S5: Fig. S1).

### 3.4. Scope and broader implications

Temporal root dynamics and the balance between gains and losses likely differ across systems, which may influence how temporal variation is represented depending on when and how frequently observations are made. As a result, the relationship between inverse CV estimates derived from metrics that approximate pools or fluxes, as well as the extent to which these estimates depend on measurement frequency and timing, may vary across ecological contexts. Metric choice, measurement frequency, and timing should therefore be understood as defining the context in which inverse CV is quantified, rather than as inherently more or less appropriate approaches. Our study is based on a single experimental system and therefore represents a specific case of temporal root dynamics in a temperate dry, acidic grassland under low-intensity management, where aboveground biomass is mown once per year (Alonso-Crespo et al., 2025). Such systems are generally nutrient poor (Jentsch & Beyschlag, 2003), develop slowly, and have low biomass turnover (Beyschlag et al., 2009; Tibbett et al., 2019). Systems like these are likely to have positive directional consistency and relatively low pool invariability. Other systems with faster turnover, stronger seasonal replacement, or more intensive management may show higher pool invariability and directional consistency closer to zero.

Importantly, these methodological considerations are not limited to minirhizotrons but likely apply more broadly to ecological time series, including remotely sensed data. Here, results may vary among vegetation indices, sensors, and data-processing approaches (Yan et al., 2025; Zeng et al., 2020), and measurement frequency and timing are constrained by sensor characteristics and revisit intervals, which influence how well vegetation dynamics are resolved (Cui et al., 2020; Tian et al., 2021). Approaches also differ in how vegetation dynamics are summarized, including annual maximum NDVI, which represents a snapshot in time, and change-based metrics such as NDVI differences or greening and browning rates, which represent change through time (Kumar et al., 2022; Reed et al., 1994; N. Shan et al., 2025). As a result, inverse CV estimates derived from remotely sensed vegetation indices are likewise expected to depend on how ecosystem dynamics are represented, as well as when and how frequently they are observed.

Similar considerations arise for other ecosystem fluxes, including CO₂, CH₄, and sensible heat, which can take both positive and negative values. Despite the known difficulties of applying the CV to such variables (Kvålseth, 2017), studies have used it to quantify variability in these fluxes (Evrendilek et al., 2011; Peltola et al., 2015; W. Zhang et al., 2018). When the mean approaches zero, the CV (σ/μ) increases in magnitude, whereas the inverse CV (μ/σ) approaches zero, provided temporal variability is non-zero. For the inverse CV, the sign retains information about the prevailing direction of the flux, while the absolute magnitude reflects the strength of the directional signal over time. Together, these properties provide a basis for interpreting the inverse CV for signed fluxes in terms of directional consistency.

### 3.5. Recommendations and conclusions

Our results show that inverse CV estimates from minirhizotron observations depend on how root dynamics are represented, as well as when and how frequently they are measured. These findings have important implications for how inverse CV is applied and interpreted. Overall, because abundance– and change-based metrics capture different dimensions of temporal variation and can lead to divergent interpretations, the choice of the underlying metric should be aligned with the ecological question. Standing root abundance can be used to characterize invariability of the standing root pool, whereas change-based metrics characterize directional consistency in root dynamics. For minirhizotron observations, where both abundance– and change-based metrics can be derived from the same time series, reporting both may provide complementary insight into root dynamics. Similarly, because measurement frequency and timing also influence the variation detected in the time series and resulting inverse CV estimates, they should be explicitly reported and justified.

Comparisons between inverse CV above– and belowground require particular care because the underlying measurements are not always conceptually aligned. When above– and belowground dynamics are quantified using comparable metrics, direct comparisons between realms may be appropriate. This could be the case in forest systems, where pool– and flux-based measurements aboveground can be matched with corresponding measures derived from minirhizotron observations. When direct comparisons are made, measurement frequency and timing should be matched. In grasslands, the conceptual alignment is less straightforward. When using minirhizotrons, annual peak standing root abundance may provide the closest comparison with annually harvested aboveground biomass. However, peak root abundance was identified retrospectively and its timing varied among years and tubes, making it not fully equivalent to harvesting aboveground biomass near an expected community-level peak. For measurements that are even less directly comparable, it may be more appropriate to evaluate above– and belowground inverse CV in parallel rather than comparing estimates directly. Finally, because differences in the representation as well as measurement frequency and timing of ecosystem dynamics can lead to non-equivalent estimates, comparisons of inverse CV estimates across studies should be made with caution.

#### Text Box 1. Conceptual distinction between pools and fluxes

Ecosystem properties can be described in terms of quantities present at a given time (pools, such as standing biomass or carbon pools) or quantities that describe gains and losses through time (fluxes, such as productivity or decomposition), including their net balance (e.g., net biomass change). Pools and fluxes are linked (Chapin et al., 2011; Clark et al., 2001). The size of a pool at a given time is determined by the accumulated balance between gains and losses (Fig. T1). Fluxes quantify these gains and losses and thereby drive changes in pools. Despite their linkage, these quantities are not equivalent. For example, two sites can show similar biomass yet differ strongly in turnover: high production offset by high mortality can maintain a stable biomass pool, while low production and low mortality can yield a similar pool with much slower turnover (Clark et al., 2001; Finér et al., 2011; terHorst & Munguia, 2008). Distinguishing between pools and fluxes therefore clarifies whether an ecosystem function is characterized in terms of its stored quantity or the processes regulating its turnover.

The inverse CV (μ/σ) can be calculated for both pools and fluxes. However, estimates may differ even when they originate from the same underlying root dynamics (Fig. T1), because pools and fluxes capture different aspects of that system. In contrast to standing-abundance measures, net change values can be negative when root mortality outpaces growth, complicating their interpretation relative to commonly used biomass-based proxies for productivity, which are non-negative.

Empirical studies using the inverse CV often characterize it as the invariability of aboveground productivity (e.g., Hautier et al., 2015; Isbell et al., 2009). The way aboveground biomass is typically measured, however, results in a quantity that is neither strictly a pool nor a flux. For example, in grassland biodiversity experiments, the inverse CV is calculated from the annual harvest of aboveground biomass at peak biomass. This represents biomass that has accumulated over one year. It is therefore not a direct measure of productivity, because biomass produced and lost during this period is not captured. Nor is it a measure of standing biomass, because the biomass is removed annually or biannually. This can complicate comparisons with minirhizotron-derived measures, as these observations can be represented as standing root length density (a proxy for a pool) or changes in standing root length density (a proxy for a flux).

**Fig. T1.**
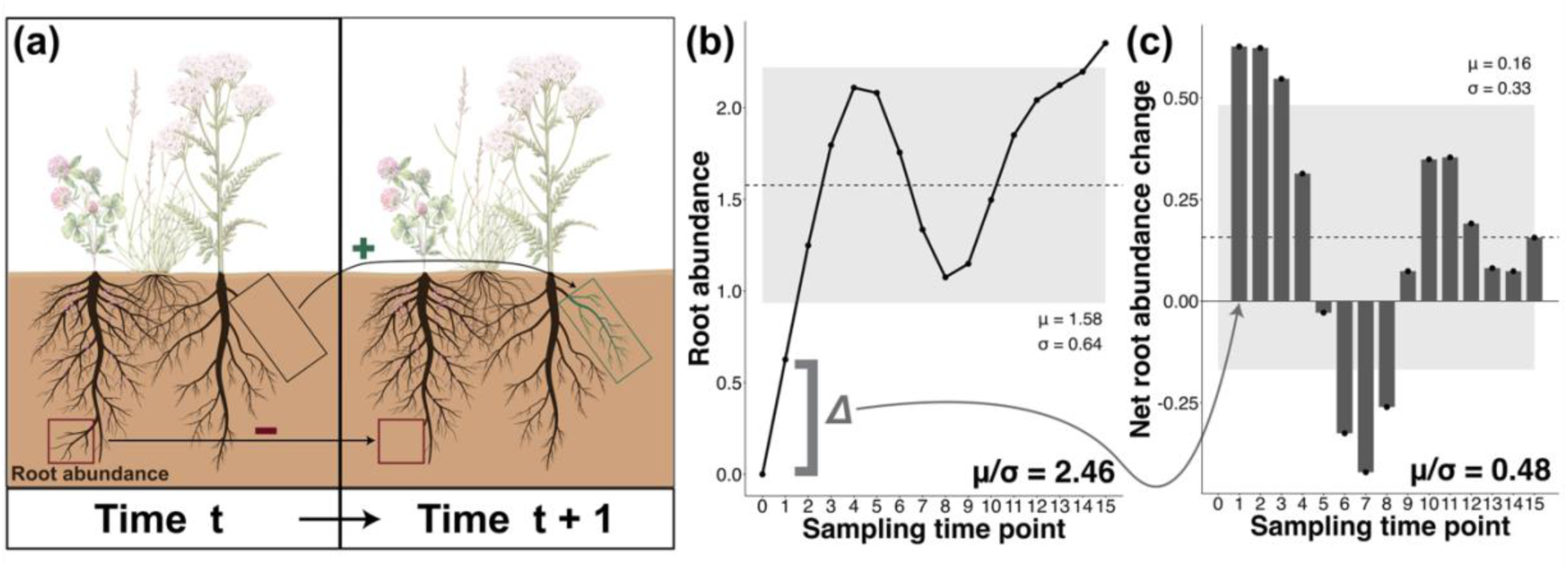
Conceptual illustration of root abundance (pool) and its net change (flux). (a) Root abundance is observed at consecutive time points (t and t+1), with roots gained (green) and lost (red) between observations. (b) When the system is expressed as root abundance, a quantity present at each time point, the resulting time series represents a pool. (c) When the system is expressed as the net change between consecutive observations, the resulting time series represents a flux. Dashed lines represent the calculated mean (μ), and the grey shaded areas represent the standard deviation (σ).

## Notes

### Competing Interest Statement

The authors have declared no competing interest.

### Funding

- Deutsche Forschungsgemeinschaft: 420444099
- NWO – Exacte en Natuurwetenschappen: VI.Vidi.213.052

