## Appendix S1 for "Quantifying and interpreting the inverse coefficient of variation from minirhizotron-derived root dynamics"

### Appendix S1. Bayesian model validation

**Table S1** Leave-one-out cross-validation (LOO-CV) comparison of alternative hierarchical structures for the Bayesian distributional model. Differences in expected log predictive density (ΔELPD) and their standard errors are reported relative to the model with the highest ELPD. Tube identifiers were unique across plots and plant functional group order-of-arrival treatments. Additional hierarchical effects did not clearly improve predictive performance, so we retained the simplest supported hierarchical structure.

| Hierarchical structure | ΔELPD | SE(ΔELPD) |
|---|---|---|
| Tube | 0.0 | 0.0 |
| Plot / Tube | -18.2 | 17.9 |
| Arrival / Plot / Tube | -18.9 | 18.4 |

**Table S2** Summary table of the BRMS model results. The table summarizes the posterior distributions of model parameters grouped by model component. Reported values include the posterior means, standard errors, 95% credible intervals (CI), R-hat values, and effective sample sizes (Bulk ESS and Tail ESS) for each parameter.

| **Parameter** | **Estimate** | **Est. error** | **Lower CI** | **Upper CI** | **R-hat** | **Bulk ESS** | **Tail ESS** |
|---|---|---|---|---|---|---|---|
| **Measurement frequency and timing** | | | | | | | |
| All measurements | 1.57 | 0.03 | 1.51 | 1.63 | 1.00 | 4719 | 7063 |
| Monthly measurements | 1.69 | 0.03 | 1.63 | 1.75 | 1.00 | 8750 | 8263 |
| Pre-harvest measurements | 1.82 | 0.25 | 1.32 | 2.33 | 1.00 | 6431 | 6669 |
| Peak measurements | 2.19 | 0.15 | 1.91 | 2.48 | 1.00 | 5809 | 6270 |
| **Metric** | | | | | | | |
| Net change | -1.37 | 0.03 | -1.43 | -1.31 | 1.00 | 5530 | 7351 |
| Rate of change | -1.32 | 0.03 | -1.37 | -1.26 | 1.00 | 5532 | 7553 |
| **Measurement frequency and timing × metric** | | | | | | | |
| Monthly measurements × Net change | -0.09 | 0.04 | -0.17 | -0.01 | 1.00 | 5601 | 7188 |
| Pre-harvest measurements × Net change | 0.62 | 0.26 | 0.09 | 1.14 | 1.00 | 6443 | 7012 |
| Peak measurements × Net change | 0.22 | 0.15 | -0.09 | 0.52 | 1.00 | 5598 | 6264 |
| Monthly measurements × Rate of change | -0.12 | 0.04 | -0.20 | -0.03 | 1.00 | 5815 | 7875 |
| Pre-harvest measurements × Rate of change | 0.70 | 0.27 | 0.15 | 1.24 | 1.00 | 6456 | 7556 |
| Peak measurements × Rate of change | 0.14 | 0.16 | -0.17 | 0.46 | 1.00 | 5866 | 6796 |
| **Residual variance model** | | | | | | | |
| Residual SD: All measurements | -1.75 | 0.08 | -1.90 | -1.59 | 1.00 | 10388 | 9018 |
| Residual SD: Monthly measurements | -1.75 | 0.08 | -1.90 | -1.60 | 1.00 | 11204 | 9207 |
| Residual SD: Pre-harvest measurements | 0.45 | 0.14 | 0.17 | 0.72 | 1.00 | 3572 | 6613 |
| Residual SD: Peak measurements | -0.12 | 0.14 | -0.39 | 0.15 | 1.00 | 3632 | 5960 |
| Residual SD: Net change | -1.16 | 0.14 | -1.43 | -0.88 | 1.00 | 3143 | 5772 |
| Residual SD: Rate of change | -0.86 | 0.13 | -1.10 | -0.60 | 1.00 | 3578 | 6584 |
| **Hierarchical parameter** | | | | | | | |
| sd(Intercept) | 0.06 | 0.01 | 0.04 | 0.08 | 1.00 | 3538 | 5277 |

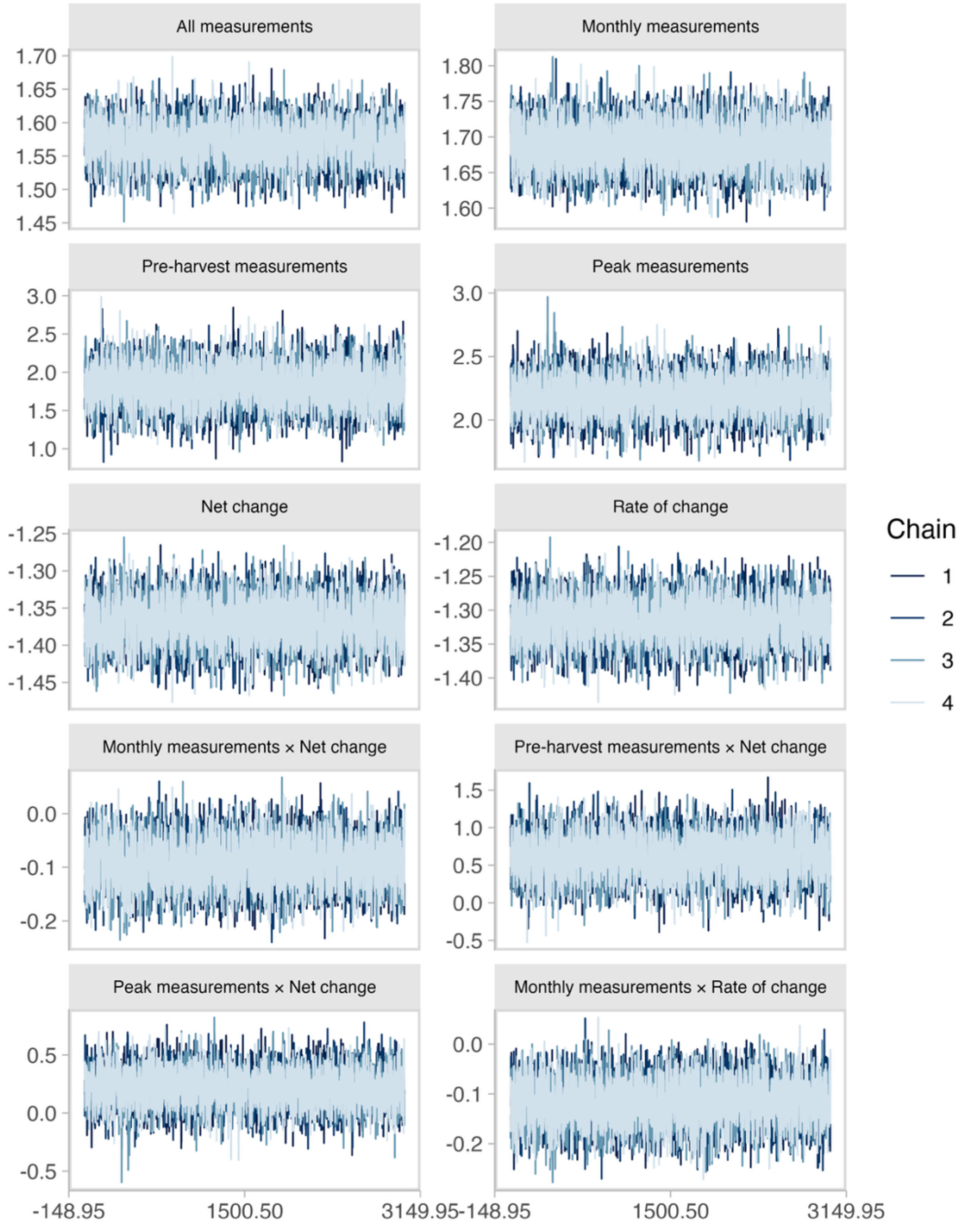


**Fig. S1** Trace plots of MCMC chains for all model parameters, used to assess convergence and mixing across chains.

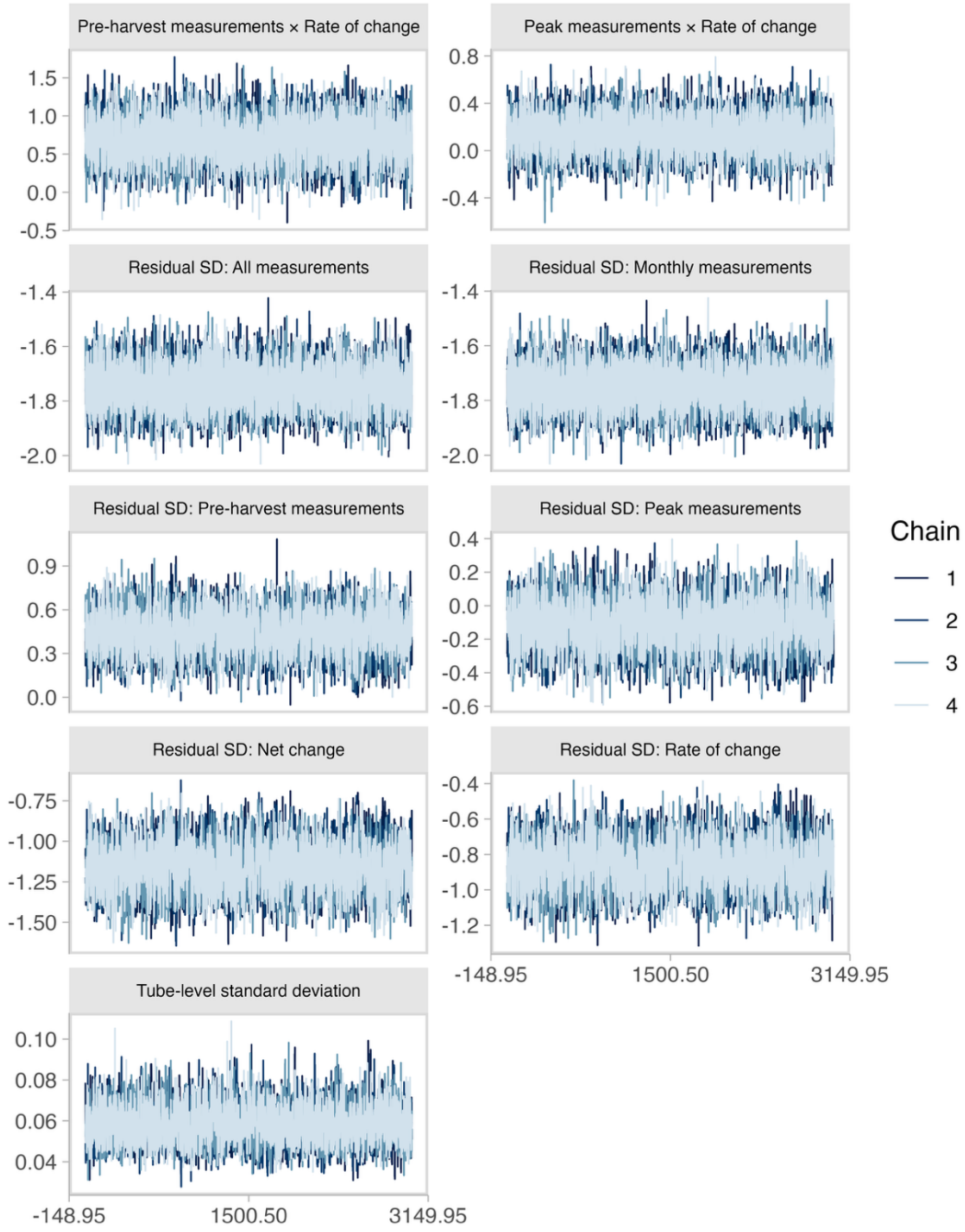


**Fig. S1 (continued).** Trace plots of MCMC chains for all model parameters, used to assess convergence and mixing across chains.

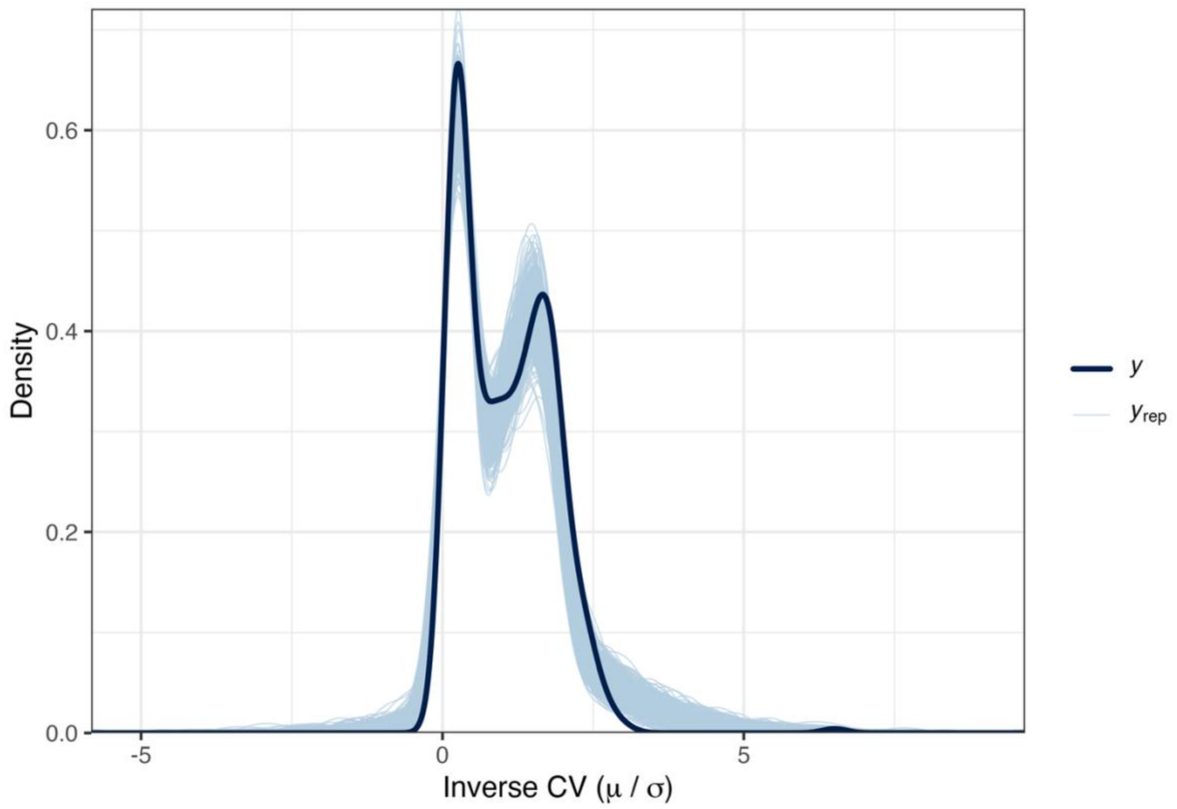


**Fig. S2** Posterior predictive check based on density overlay. The observed data (*y*, dark blue line) are compared to the fitted draws of the models (*y_rep*, light blue lines). Close agreement indicates that the model captures the overall distribution of the data.

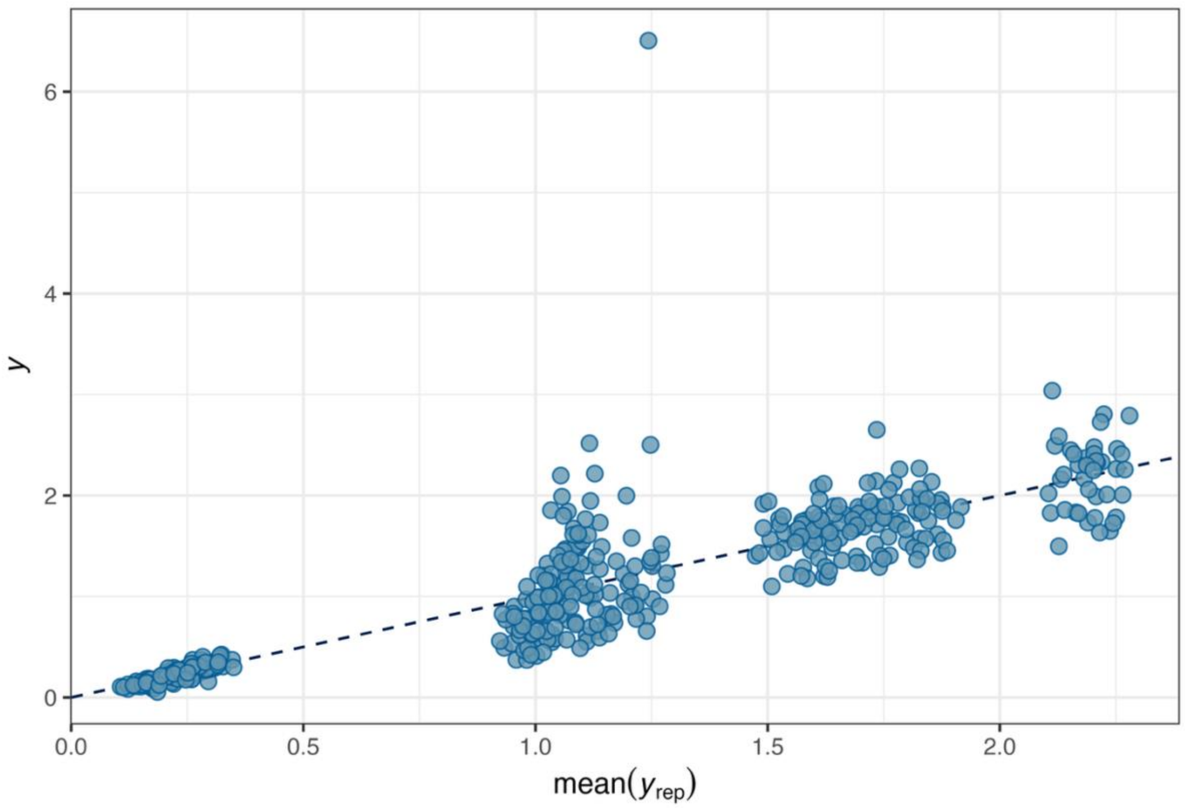


**Fig. S3** Posterior predictive check based on average predictions. Observed values (*y*) are compared to the mean of the posterior predictive distribution (*y_rep*). Agreement along the 1:1 line indicates that the model adequately reproduces the central tendency of the data.

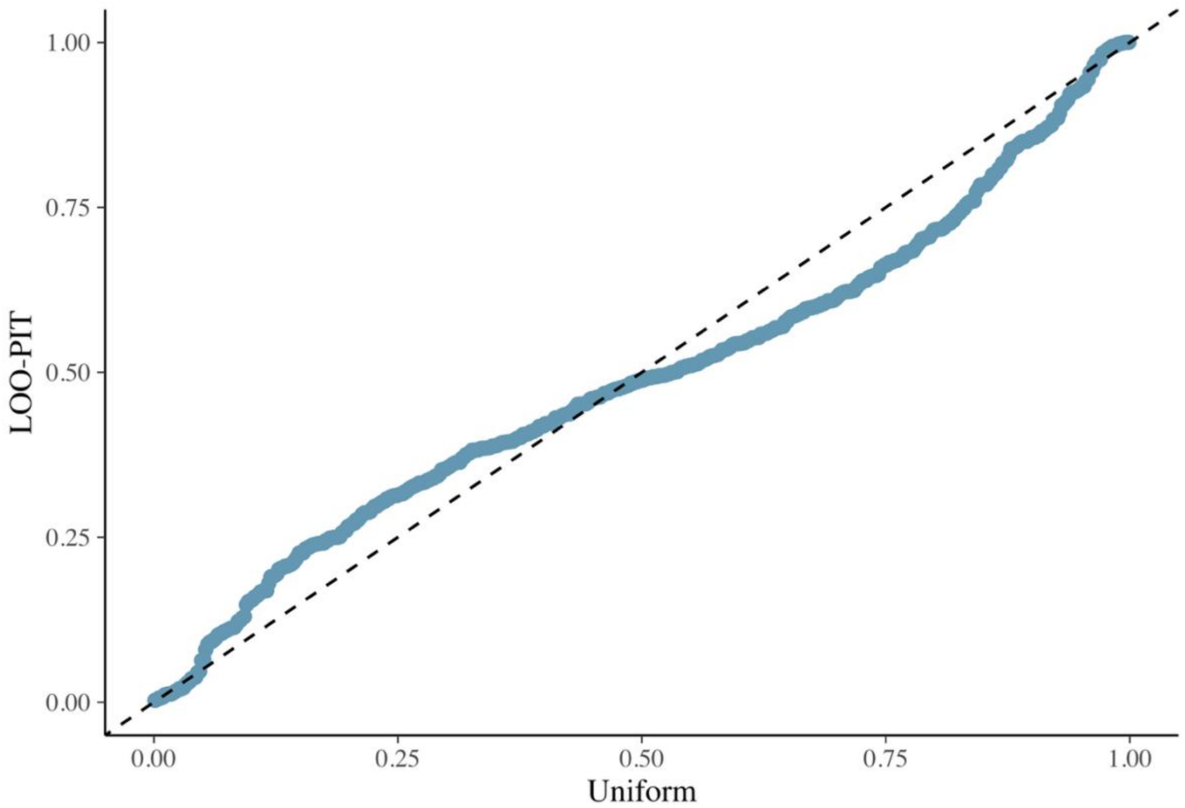


**Fig. S4** LOO-PIT (Leave-One-Out Probability Integral Transform) calibration. The empirical cumulative distribution of LOO-PIT values (solid blue line) is compared to the expected uniform distribution (dashed diagonal line).

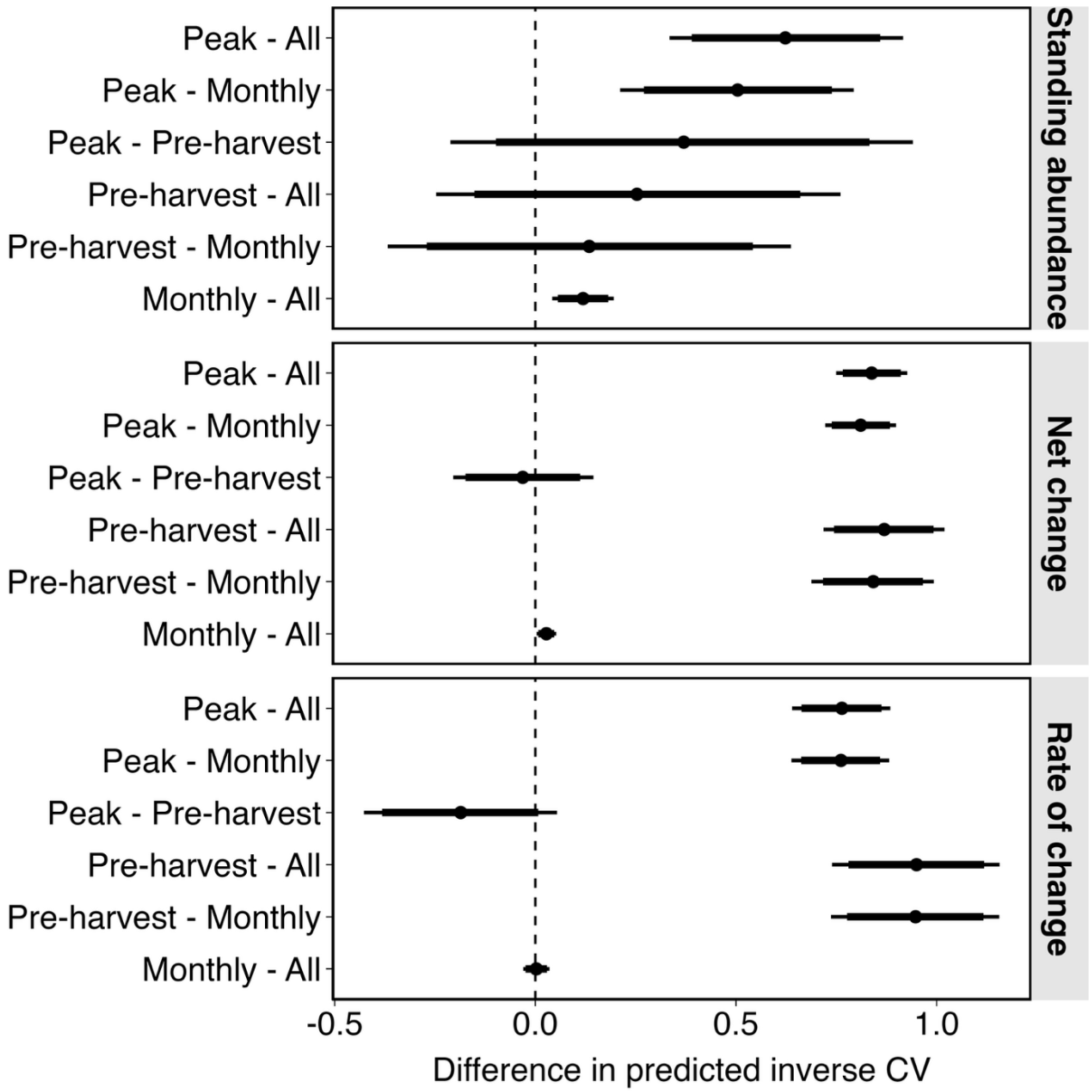


**Fig. S5** Pairwise contrasts among measurement frequency and timing within each metric. Points indicate mean differences in predicted inverse CV, and bars show 89% (thick) and 95% (thin) credible intervals.
