## Appendix S2 for "Quantifying and interpreting the inverse coefficient of variation from minirhizotron-derived root dynamics"

### Appendix S2. GAM-based approach

In addition to the raw-data approach described in the main text, we implemented a complementary GAM-based approach that models smoothed temporal trajectories of *pRLD*. From these smooth functions, rates of change (first derivatives) and net change (integrals) between consecutive sessions can be derived.

#### 1. Deriving GAM-based metrics

##### 1.1. Model specification

In the GAM-based approach, we modeled temporal changes in *pRLD* using generalized additive models (GAMs). For each of the four time series representing the different measurement frequencies and timings, we fitted a single model that included all tubes. The response variable was depth-averaged *pRLD*, and time (days since the start of POEM2021) was treated as a continuous predictor. To retain the same data structure as in the raw-data approach presented in the main text and allow each tube to follow its own temporal trajectory, we implemented tube-specific smooths using a factor–smooth interaction, which provides a separate smooth for each tube while sharing a common smoothing penalty. Models were fitted using the *mgcv* package (Wood, 2017) in R (R Core Team, 2025), with the basis dimension ($k$) set to the number of available time points for each measurement frequency, allowing sufficient flexibility while preventing overfitting through penalization. Smoothing parameters were estimated via restricted maximum likelihood (REML). We used a Tweedie distribution with a log link to accommodate the non-negative, continuous nature of *pRLD* measurements and the presence of zeros.

##### 1.2. Derivation of abundance and change metrics

Standing root abundance was quantified from the fitted model for each of the four time series. For each tube and session, predicted depth-averaged *pRLD* served as a proxy for standing root abundance, analogous to the raw *pRLD* values used in the raw-data approach.

Changes in root abundance were derived from the GAM-fitted curves. We quantified instantaneous rates of change in root abundance as the first derivative of the predicted *pRLD* curve with respect to time using the *gratia* R package (Simpson & Singmann, 2022). This metric is conceptually comparable to the time-standardized rate of change (*ΔpRLD / Δt*) derived from the raw data. Net change in root abundance between consecutive imaging sessions was quantified by integrating the derivative over each interval, yielding the model-implied difference in *pRLD* between those time points, comparable to *ΔpRLD* from the raw-data approach described in the main text. As in the raw-data approach, positive values indicate net root gain and negative values indicate net root loss resulting from mortality exceeding growth.

#### 2. Calculating the inverse coefficient of variation

As in the main text, inverse CV (μ/σ) was computed separately for the three metrics (predicted standing root abundance, derivative-based rate of change, and integral-based net change) within each of the four tube-specific time series (all measurements, monthly measurements, annual pre-harvest measurements, and annual peak measurements). This resulted in 12 inverse CV estimates per tube.

#### 3. Statistical analysis

GAM-derived inverse CV estimates were analyzed using the same statistical models described in the main text, with the same model structure and estimation procedures. The only difference was the use of GAM-derived rather than raw-data derived inverse CV estimates.

#### 4. Results

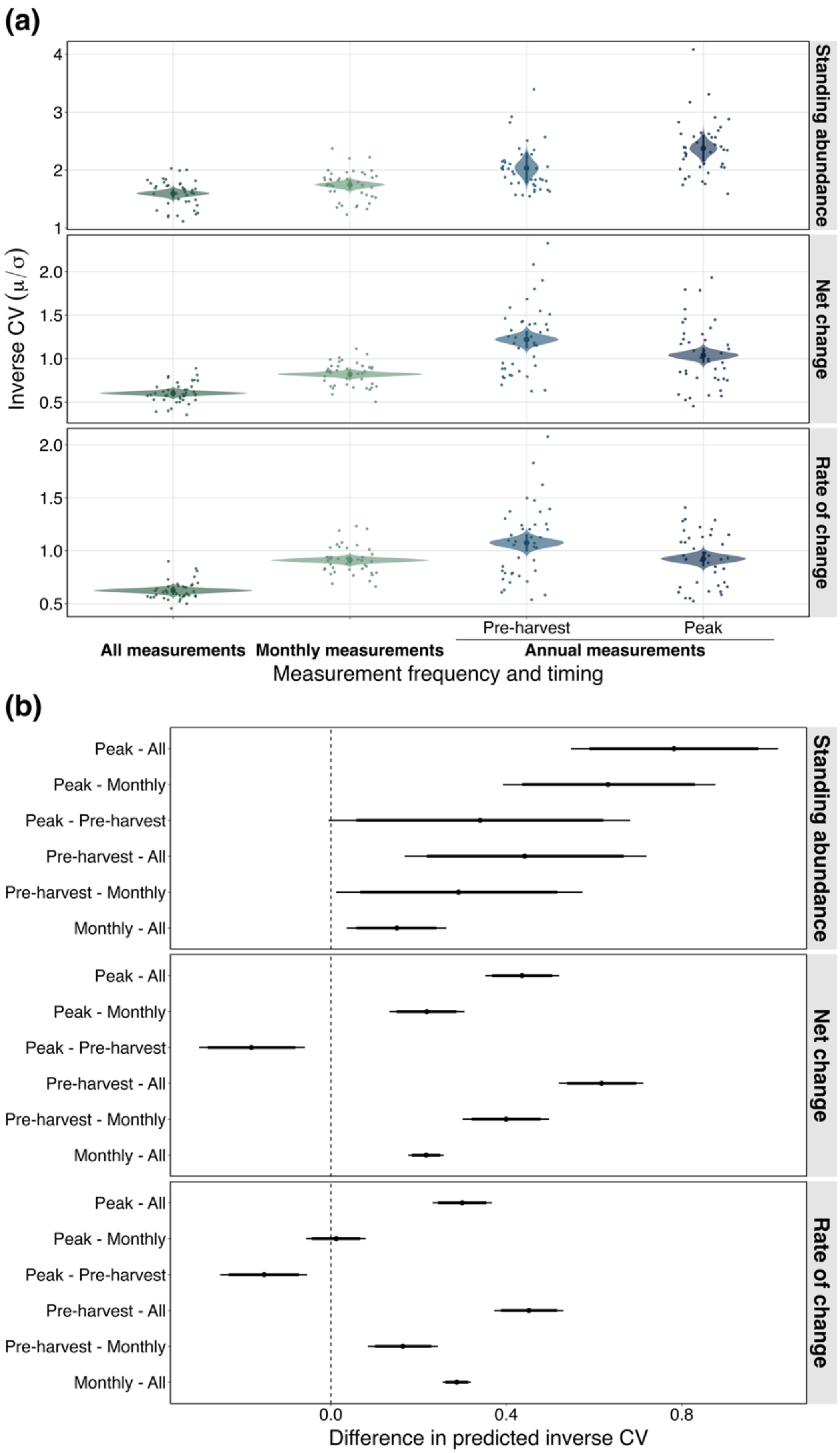
(a)
Standing abundance
Net change
Rate of change
Inverse CV (μ/σ)
Pre-harvest
Peak
All measurements
Monthly measurements
Annual measurements
Measurement frequency and timing
(b)
Peak - All
Peak - Monthly
Peak - Pre-harvest
Pre-harvest - All
Pre-harvest - Monthly
Monthly - All
Standing abundance
Net change
Rate of change
Difference in predicted inverse CV

**Fig. S1** Effects of measurement frequency and timing, and metric on GAM-derived inverse CV estimates. (a) Inverse CV (μ/σ) calculated from GAM-derived standing root abundance (predicted pRLD), integral-based net change, and derivative-based rate of change for all, monthly, and annual measurements at two timings (pre-harvest and peak). Points show tube-level GAM-derived estimates; model summaries indicate posterior means with 95% credible intervals. (b) Pairwise contrasts in predicted inverse CV among measurement frequencies and timings for each metric. Points denote posterior mean differences, and bars represent 89% (thick) and 95% (thin) credible intervals. Intervals excluding zero indicate credible differences. For standing root abundance, inverse CV increased with coarser measurement frequency and was higher for annual peak than annual pre-harvest measurements. Change-based metrics showed non-monotonic responses that differed between net change and rate of change. For net change, inverse CV increased from all to pre-harvest measurements and decreased again at peak measurements, whereas for rate of change, monthly, pre-harvest, and peak measurements were all higher than all measurements, with monthly and peak measurements showing similar inverse CV and both remaining lower than pre-harvest measurements.
