## Appendix S3 for "Quantifying and interpreting the inverse coefficient of variation from minirhizotron-derived root dynamics"

### Appendix S3. Information-based stability

In addition to the inverse coefficient of variation (*inverse CV* = $\mu/\sigma$) presented in the main text, we quantified the invariability of standing root abundance using the information-based framework proposed by Chao et al. (2025). This framework is defined for non-negative data and is parameterized by an order (q) that determines sensitivity to the magnitude of values in the time series. Here, we considered q = 1 and q = 2, with q = 2 being more sensitive to larger values.

#### 1. Calculation of information-based stability

Information-based stability was computed for depth-averaged *pRLD* time series using the iSTAY_Single() function from the *iSTAY* R package (Chao et al., 2025) for each tube across three measurement frequencies and two annual measurement timings, resulting in four time series (all measurements, monthly measurements, annual pre-harvest measurements, and annual peak measurements). Information-based stability was calculated separately for $q = 1$ and $q = 2$.

#### 2. Statistical analysis

Information-based stability estimates were analyzed using Bayesian hierarchical models implemented in brms. For each order ($q = 1$ and $q = 2$), information-based stability was modeled as a function of measurement frequency and timing, with tube included as a hierarchical grouping variable to account for non-independence among estimates derived from the same tube across the four time series.

The models were fitted without an intercept and with default weakly informative priors using four Markov Chain Monte Carlo chains with 4,000 iterations each, including 1,000 warm-up iterations. The models showed excellent convergence (all R-hat = 1.00), and trace plots indicated adequate mixing of chains. Model performance and goodness-of-fit were evaluated using Leave-One-Out Cross-Validation and posterior predictive checks. Pairwise contrasts among measurement frequencies and timings were computed from posterior draws to quantify differences in predicted information-based stability.

#### 3. Results

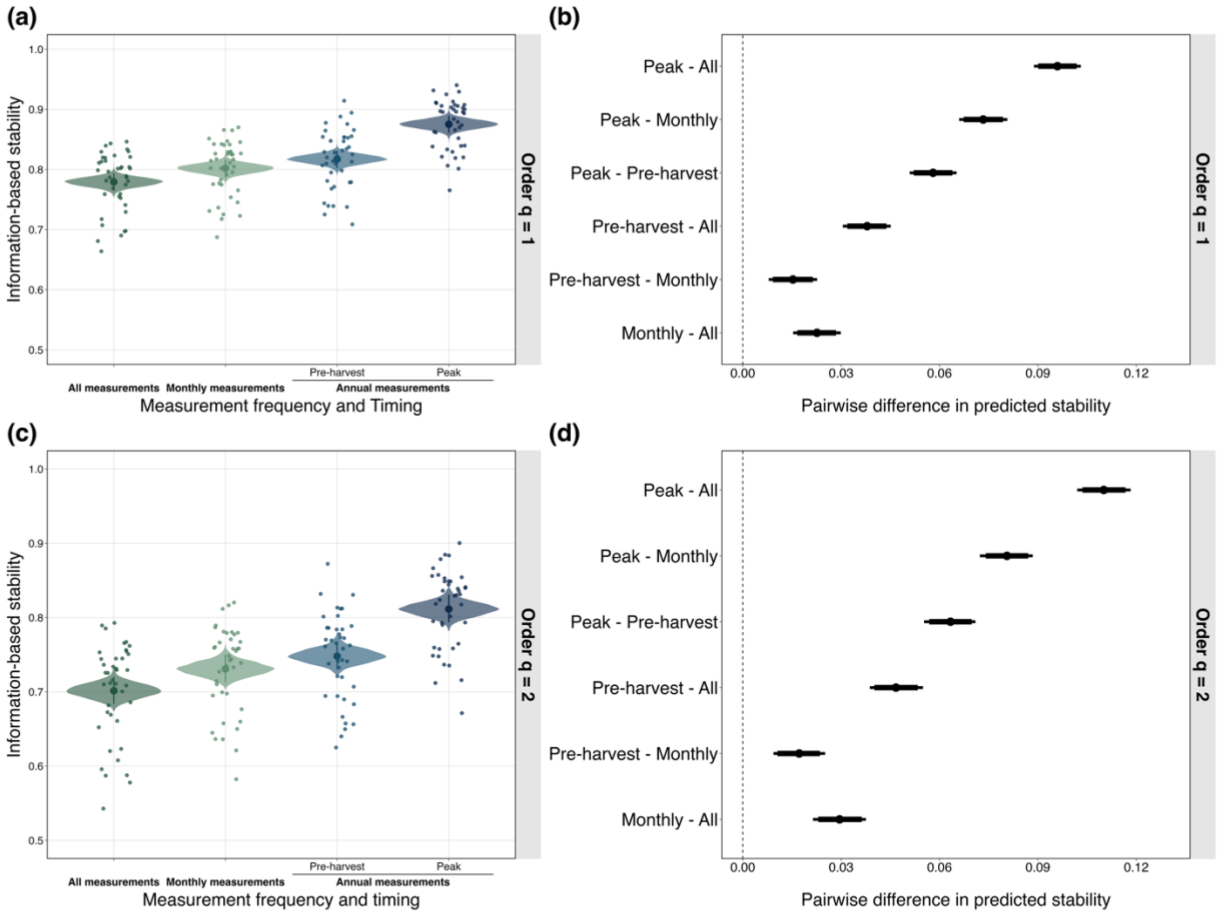


**Fig. S1** Information-based stability of standing root abundance (*pRLD*) and corresponding pairwise contrasts across four time series representing three measurement frequencies and two annual measurement timings (all measurements, monthly measurements, annual pre-harvest measurements, and annual peak measurements). (a, c) Information-based stability for q = 1 (a) and q = 2 (c). Points represent observed tube-level estimates, and intervals show posterior means with 95% credible intervals from Bayesian hierarchical models. (b, d) Pairwise contrasts in predicted information-based stability among measurement frequencies and timings for q = 1 (b) and q = 2 (d). Points denote posterior means; bars represent 95% credible intervals. Information-based stability differed among all four time series, with higher estimates at coarser measurement frequencies and for annual peak than annual pre-harvest measurements (all 95% credible intervals for pairwise contrasts excluded zero).
