## Appendix S4 for "Quantifying and interpreting the inverse coefficient of variation from minirhizotron-derived root dynamics"

### Appendix S4. Conceptual examples illustrating the distinction between pool invariability and directional consistency

The ecological meaning of the inverse coefficient of variation (inverse CV = $\mu/\sigma$) depends on whether it is calculated from a pool (e.g., standing root abundance) or from a flux (e.g., change in root abundance between successive observations). Here, we use conceptual examples to illustrate this distinction for root abundance and net change in root abundance. Because all time steps in these examples are equal to one, net change is equivalent to rate of change, as dividing net change by the length of the observation interval does not change the values.

When the inverse CV is calculated from root abundance, it reflects the invariability of a maintained standing root pool. The mean ($\mu$) represents average root abundance across observations, whereas the standard deviation ($\sigma$) represents temporal variation in abundance around that mean. A high inverse CV therefore indicates that temporal variation is small relative to mean root abundance, whereas a low inverse CV indicates greater temporal variation relative to the temporal mean. Because root abundance reflects the cumulative outcome of gains and losses, their temporal ordering can affect both the mean abundance and its variation through time (Fig. S8). A relatively invariant pool can therefore be maintained when gains and losses are offset through time, so that biomass fluctuates around a maintained level. In such cases, root abundance can appear relatively stable at the pool level even though turnover occurs through largely alternating gains and losses (Fig. S8 c,d; Fig. S9 a,b).

When the inverse CV is calculated from change-based metrics, its interpretation shifts because losses can outpace gains over time, resulting in a negative mean and therefore a negative inverse CV. This complicates an interpretation in terms of invariability because the inverse CV can be positive or negative, with its sign reflecting the direction of change. In this case, inverse CV expresses mean change in standing root abundance relative to its temporal variability, with the signed mean indicating the prevailing direction of change toward gains or losses, and variability among increments contributing to how consistently that direction is reflected over time. For a given mean change, lower variability among increments therefore indicates greater consistency in the direction of change over time. We therefore interpret inverse CV calculated from change-based metrics as directional consistency, where the sign indicates the prevailing direction toward gains or losses, the absolute magnitude reflects the strength of the mean direction relative to variability among changes over time, and an inverse CV of zero indicates no prevailing direction of change. Strictly, $\mu/\sigma$ is undefined when temporal variability is zero ($\sigma = 0$), which would require identical change values across all observations. Such a case is unlikely in temporally varying ecological data.

Importantly, the temporal ordering of change increments does not affect their mean, standard deviation, or directional consistency, such that the same directional consistency can arise from different temporal sequences of gains and losses (Fig. S8 b,d). When gains and losses are relatively balanced across the observed period, mean net change is small and directional consistency is therefore low. As mean net change approaches zero, the inverse CV also approaches zero, and variability among increments has progressively less influence on the resulting inverse CV. This does not alter its interpretation as directional consistency, because a mean close to zero itself indicates little prevailing direction toward either gains or losses. Similarly, alternating gains and losses can result in low directional consistency when they produce little net change over time (Fig. S9 b). In both cases, there is little to no prevailing direction in root gains or losses over the observed period. In contrast, under sustained growth, successive net gains can be similar in magnitude, resulting in a positive mean net change with relatively little variation among increments and therefore greater directional consistency (Fig. S9 d). Yet, sustained growth results in lower pool invariability than when root abundance fluctuates around a maintained level (Fig. S9 a,c).

Disturbances that result in higher-than-average root loss may modify pool invariability and directional consistency in similar ways. For example, root loss may become more pronounced under drought, shifting the balance between gains and losses across time. As a consequence, both pool invariability and directional consistency may decrease relative to control conditions (Fig. S10). However, the underlying drivers can differ. In the conceptual scenario illustrated here, the reduction in pool invariability arises primarily from a lower mean abundance, whereas its variability changes only marginally (Fig. S10 a,c). By contrast, the decrease in directional consistency reflects both a shift in mean net change and an increase in variability among increments (Fig. S10 b,d). Although the abundance- and change-based inverse CVs shift in the same direction, they may do so for different reasons. Together, these examples illustrate that pool invariability and directional consistency capture distinct dimensions of temporal variation, reflecting variation in the state of a pool and the overall tendency toward gains or losses through time, respectively.

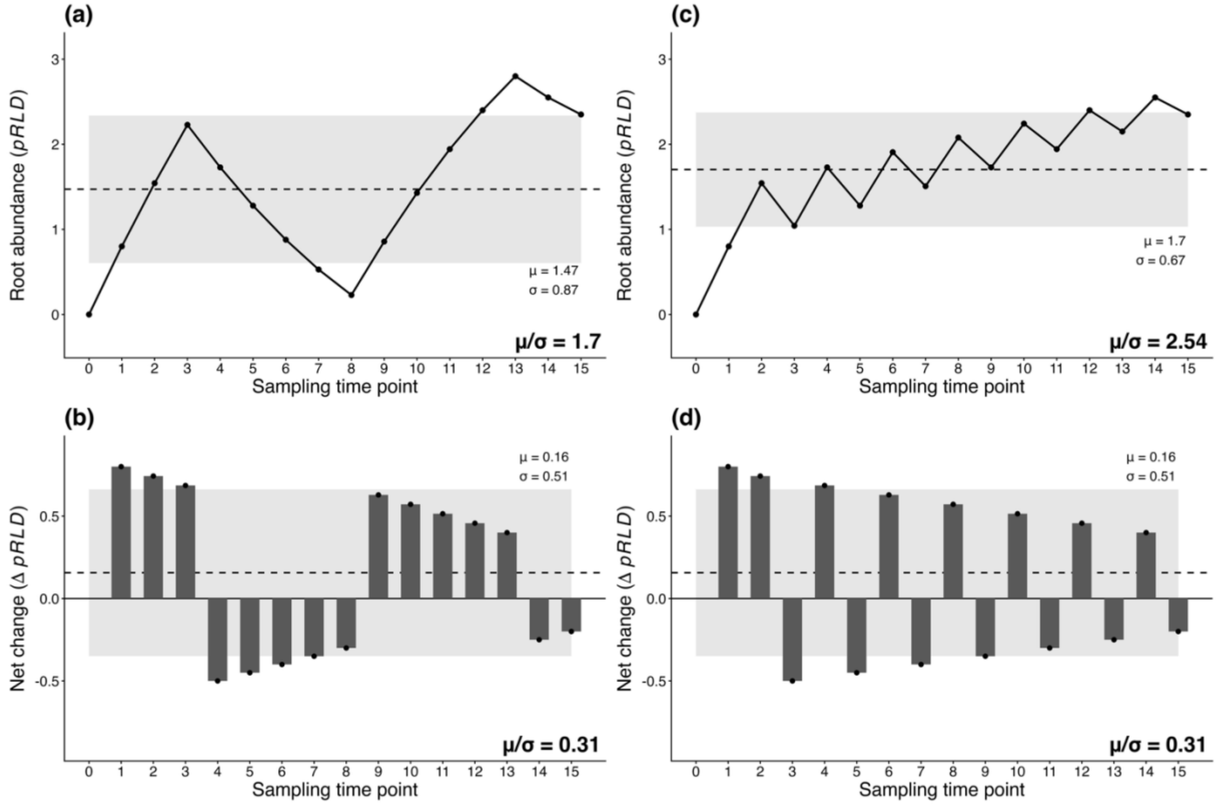


**Fig. S1** Conceptual illustration of how the temporal ordering of net-change increments may affect pool invariability but not directional consistency. Panels a-b show root abundance (*pRLD*) through time (a) and the corresponding net change between consecutive observations (*ΔpRLD*; b) when gains and losses occur in sequential runs. Panels c-d show the same set of net-change increments arranged so that gains and losses largely alternate through time, again shown as root abundance (c) and net change (d). Because the net-change increments are identical and differ only in their temporal ordering, their mean, standard deviation, and directional consistency (μ/σ) are identical in panels b and d, whereas both the mean and temporal variation in root abundance, and hence root pool invariability (μ/σ), differ between panels a and c. Shaded areas indicate ±1 standard deviation around the mean (dashed line).

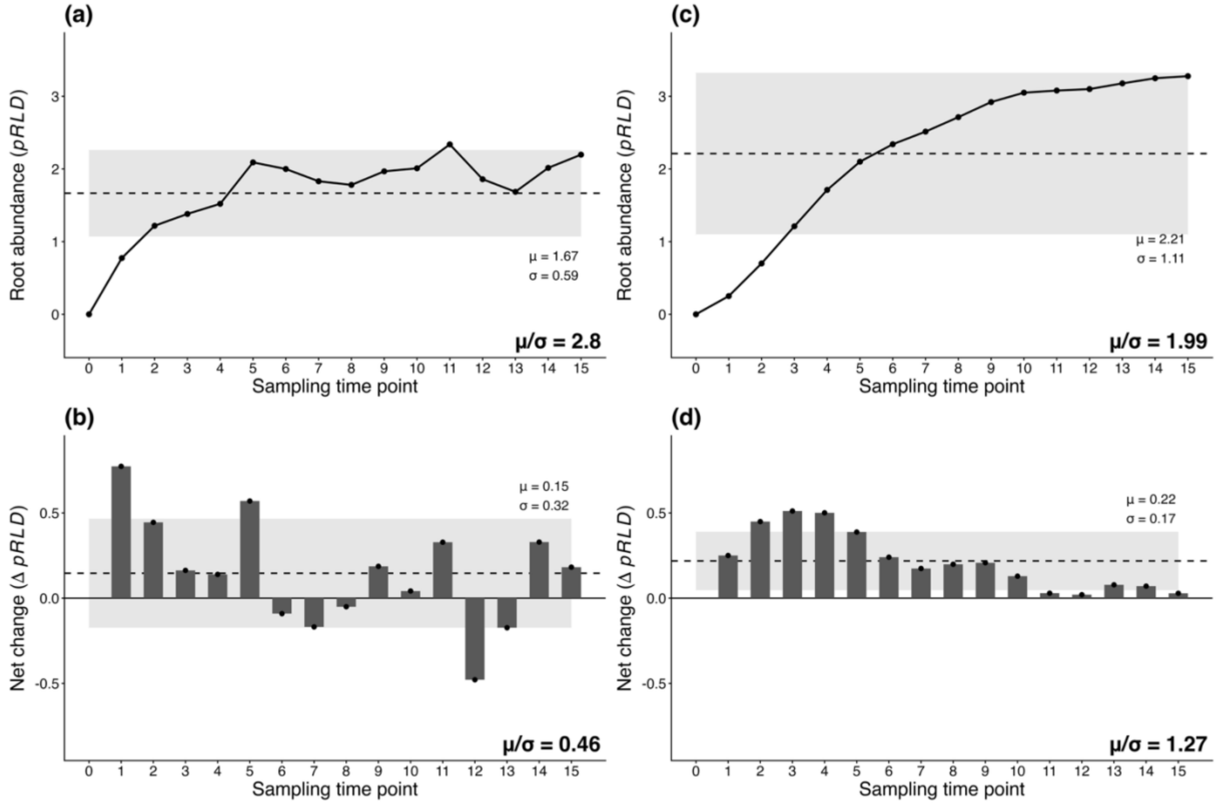


**Fig. S2** Conceptual figure illustrating that pool invariability and directional consistency quantify different aspects of temporal variation in root abundance. Panels a-b represent an equilibrium scenario, shown as root abundance (*pRLD*) through time (a) and the corresponding net change between consecutive observations (*ΔpRLD*; b). Panels c-d represent a scenario of sustained directional growth, again shown as abundance (c) and the corresponding net change (d). Shaded areas indicate ±1 standard deviation around the mean (dashed line).

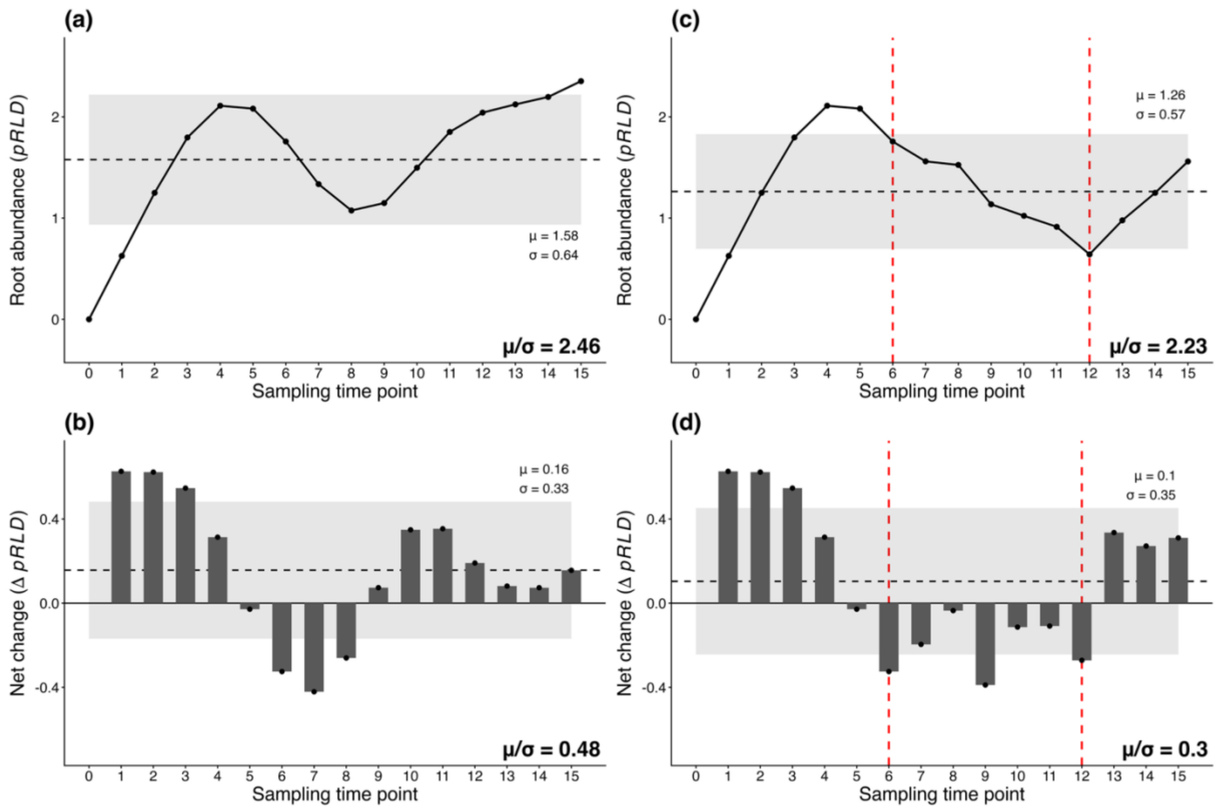


**Fig. S3** Conceptual illustration how drought may affect pool invariability and directional consistency. Panels a-b show a control scenario, and panels c-d show a drought scenario affecting root dynamics (red dashed lines). Panels a and c display root abundance (*pRLD*) through time, whereas panels b and d show the corresponding net change (*ΔpRLD*). Shaded areas indicate ±1 standard deviation around the mean (dashed line).
