## Appendix S5 for "Quantifying and interpreting the inverse coefficient of variation from minirhizotron-derived root dynamics"

### Appendix S5. Example time series from a single minirhizotron tube for illustrative purposes

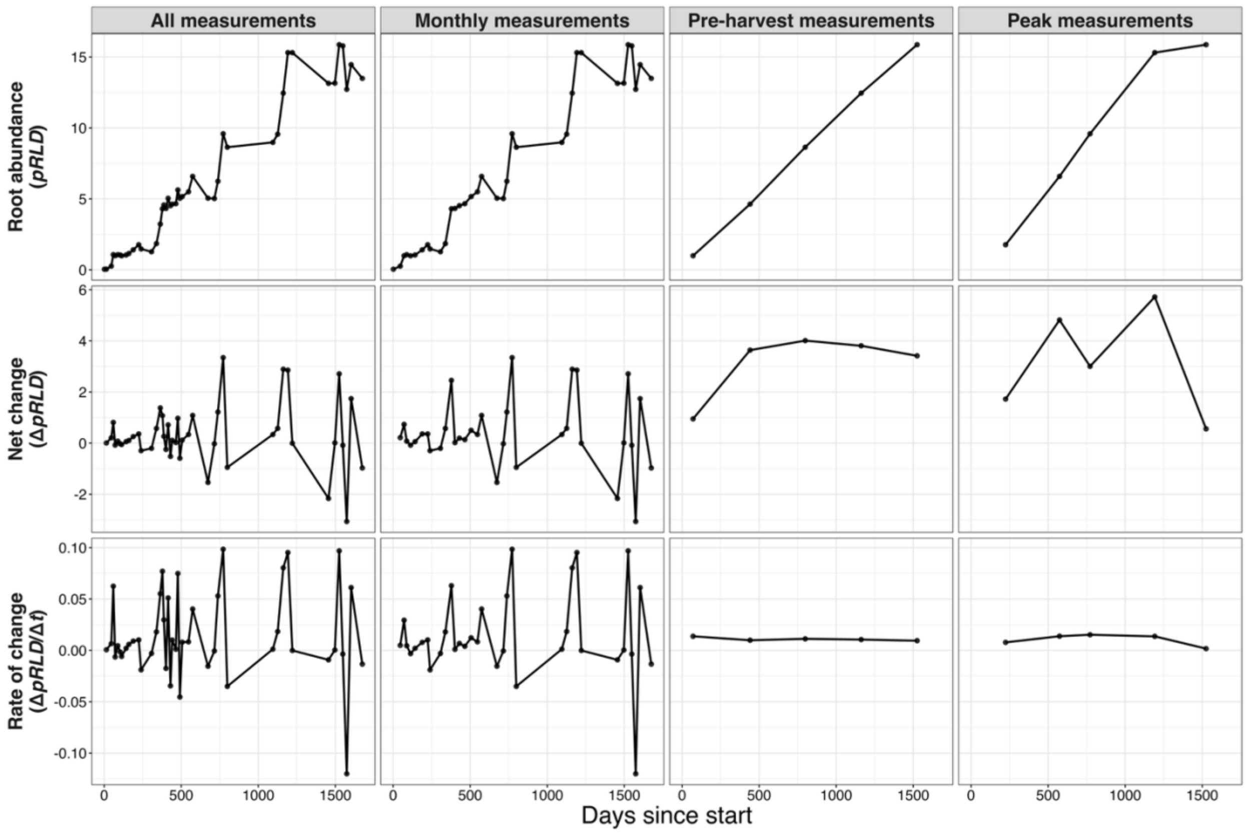


**Fig. S1** Example time series from Tube 2, located in Plot 201, illustrating standing root abundance and change-based metrics across measurement frequencies and annual timings. Columns show the four time series used in the analysis, and rows show standing root abundance (*pRLD*), net change (*ΔpRLD*), and rate of change (*ΔpRLD/Δt*). At each time point, *pRLD* is averaged across all imaged depths within the tube. The x-axis shows days since the first sowing event, marking the establishment of POEM2021.
